# Differential expression of PKCα and -β in the zebrafish retina

**DOI:** 10.1101/361303

**Authors:** Marion F. Haug, Manuela Berger, Matthias Gesemann, Stephan C. F. Neuhauss

## Abstract

The retina is a complex neural circuit in which visual information is transmitted and processed from light perceiving photoreceptors to projecting retinal ganglion cells. Much of the computational power of the retina rests on signal integrating interneurons, such as bipolar cells in the outer retina. While mammals possess about 10 different bipolar cell types, zebrafish (*Danio rerio*) has at least six ON-type, seven OFF-type, and four mixed-input bipolar cells. Commercially available antibodies against bovine and human conventional protein kinase C (PKC) α and -β are frequently used as markers for retinal ON-bipolar cells in different species, despite the fact that it is not known which bipolar cell subtype(s) they actually label.

Moreover, the expression pattern of the five *prkc* genes (coding for PKC proteins) has not been systematically determined. While *prkcg* is not expressed in retinal tissue, the other four *prkc* (*prkcaa*, *prkcab*, *prkcba*, *prkcbb*) transcripts were found in different parts of the inner nuclear layer and some as well in the retinal ganglion cell layer.

Immunohistochemical analysis in adult zebrafish retina using PKCα and PKCβ antibodies showed an overlapping immunolabeling of ON-bipolar cells that are most likely of the B_ON_ s6L or RRod type and of the B_ON_ s6 type. However, comparison of transcript expression with immunolabling, implies that these antibodies are not specific for one single zebrafish conventional PKC, but rather detect a combination of PKC -α and -β variants.

## Introduction

Bipolar cells of the vertebrate retina transmit and shape the light signal from photoreceptors to projecting ganglion cells. Broadly accepted and widely used markers for bipolar cells are protein kinases α and β (PKCα, PKCβ). First used in the late 1980s [1] they soon became a popular markers for rod bipolar cells in mammals and the corresponding mixed-type ON-bipolar cell in teleosts [2–9]. These cells are presumably labelled by PKCα and/or -β [10–12]. However, there are other bipolar cell subtypes such as the smaller BON s6 type that are also labelled by this antibody in zebrafish [12]. It is currently unclear which subset of bipolar cells are in fact labelled by these antibodies.

In the mammalian retina more than ten different subtypes of ON- and OFF-bipolar cells have been identified (e.g. [13–18]. In non-mammalian vertebrates, the number of different bipolar cell types may even be higher and can exceed 20 [19–23]. The different subtypes are classified according to their morphology and their connectivity pattern within one or more sublamina of the inner plexiform layer (IPL) [19,16,18]. ON-type bipolar cells typically send their axons to the inner sublamina “b” of the IPL whereas OFF-bipolar cells stratify in the outer sublamina “a” of this layer [24–26]. Many non-mammalian vertebrates possess a set of mixed-type bipolar cells that send axons to both IPL sublaminae and functionally show both ON- and OFF-response properties [27–29]. Although some markers for specific bipolar cell subtypes exist, inter-species comparison is problematic due to the high variability in bipolar cell subtypes and differences in connection patterns. In zebrafish 17 morphologically distinct bipolar cell subtypes have been described [19]. However, recent studies considering both axonal stratification pattern and photoreceptor connectivity for the classification of bipolar cells, suggest that the number of different bipolar cell subtypes may even be as high as 33 [20]. Since PKC antibodies are commonly used to label bipolar cells, the detailed expression profile of PKCs and the specificity of these antibodies for each PKC variant are of importance.

PKCα/β belong to the group of conventional PKCs (cPKCs) that consist of the three members PKCα, -β (in two alternatively spliced isoforms I and II), and *γ* [30]. cPKCs require diacylglycerol (DAG) along with calcium and a phospholipid such as phosphatidylserine for activation [31]. They play fundamental roles in numerous signal transduction pathways and have been linked to a number of neurological diseases [32], and retinal pathologies such as diabetic retinopathy [33]. Due to the whole genome duplication event at the base of the teleost lineage (reviewed in [34], more than one gene paralog for *prkca* and *prkcb* exists in zebrafish [35]. It is not known whether these zebrafish orthologs are recognized by the commercially available antibodies and whether there is crossreactivity between the different variants. Moreover, comparative studies about *prkc* expression in the retina are missing. The aim of this study is therefore to focus on retinal *prkc* expression and to correlate the *prkc* transcript expression with the labeling of commercially used PKCα and -β antibodies.

## Materials and methods

### Fish maintenance and breeding

Adult fish (RRID:ZIRC_ZL84) were maintained under standard conditions at 26 - 28°C in a 14-hour light/10-hour dark cycle. The wild-type strain WIK was used for all experiments described here. Embryos were raised at 28°C in E3 medium (5mM NaCl, 0.17mM KCl, 0.33mM CaCl_2_, and 0.33mM MgSO_4_). They were staged according to development in days post fertilization (dpf) (Kimmel et al., 1995). 12 adult fish were used for the experiments. All larval and adult fish used in this study were fixed between 9am and 11am. The fish were euthanized using tricaine (ethyl 3-aminobenzoate methanesulfonate; Sigma-Aldrich, Buchs, Switzerland) and iced water. All animal experiments were performed in accordance with the ARVO Statement for the Use of Animals in Ophthalmic and Vision Research and were approved by the local authorities (Veterinäramt Zürich TV4206).

### (Fluorescent) *in situ* hybridization ((F)ISH)

Primers used for the generation of RNA probes were published earlier this year (Haug, Gesemann, Berger, & Neuhauss, 2018). The plasmids containing cDNA sequences of the different *prkcs* were linearized with the appropriate restriction enzymes for T7 and Sp6, and the DNA was extracted with a standard phenol/chloroform protocol using pre-spinned RNase-free Phase-Lock tubes (5 Prime, Hamburg, Germany). Linearized DNA was *in vitro* transcribed and DIG-labeled using a kit (DIG-RNA labeling kit; Roche, Rotkreuz, Switzerland), and applied on adult zebrafish retinal sections as previously described (Haug, Gesemann, Mueller, & Nehuauss, 2013) at a concentration of approximately 2 ng/μl FISH was performed as described in (Huang, Haug, Gesemann, & Neuhauss, 2012), however, fluorescent labeling was accomplished using the TSA kit #12 (Molecular Probes, Life Technologies, Zug, Switzerland). Images were taken by confocal microscopy (CLSM SP5 and TCS LSI, Leica, Heerbrugg, Switzerland), z-stacks that covered a depth of 1 – 2 μm were selected and processed using ImageJ (Version 1.49, August 2014, Java), and further processed (brightness, contrast and gamma levels of the whole image) and arranged with Adobe Photoshop (RRID:SCR_014199) and Illustrator CS5.

### Immunohistochemistry

Adult zebrafish retinal tissue was prepared similar as described in a previous publication (Haug, Gesemann, Mueller, & Neuhauss, 2013) but the fixation time was reduced to 40 min to not overfixate the tissue. Immunohistochemical labeling was performed as previously described (Fleisch, Schönthaler, Lintig, & Neuhauss, 2008) with the following modifications: Blocking solution contained 0.3% Triton X-100 (Sigma-Aldrich) instead of Tween 20 and was applied for 30 min at RT. Different primary anti-PKCα and –β antibodies were used at the following concentrations: PKCα NBP1-19273, 1:500 (Novus, Abingdon, UK); PKCα MC5, NB 200-568, 1:1000 (Novus); PKCα MC5, 1:1000 (Genetex), PKCβ1 C16, SC209, 1:1000 (Santa Cruz Biotechnology, Heidelberg, Germany). Anti-rabbit and anti-mouse Alexa 488 (1:1000; Roche) and/or Alexa 568 (1:500; Roche) were used as secondary antibodies. Slides were mounted with Mowiol-DABCO mounting medium (10% Mowiol 4-88 (Polysciences, Warrington, USA), 25% glycerol, 2.5% DABCO (1,4-diazobicyclo[2.2.2]octane, Sigma-Aldrich) in 100 mM Tris– HCl pH 8.5) and stored in darkness at 4°C. Images were taken by confocal microscopy (CLSM SP5 and TCS LSI, Leica, Heerbrugg, Switzerland) and arranged in Fig 1. Images a-a” and b-b” cover a z-stack of 4.6 and 4.8 μm, respectively, images c-c” measure 1.6 μm in depth, and images e and f are only composed of one single plane. All images were selected and processed using ImageJ (Version 1.49, August 2014, Java), and further processed (brightness, contrast and gamma levels of the whole image) and arranged using Adobe Photoshop (RRID:SCR_014199) and Illustrator CS5.

**Figure 1:**
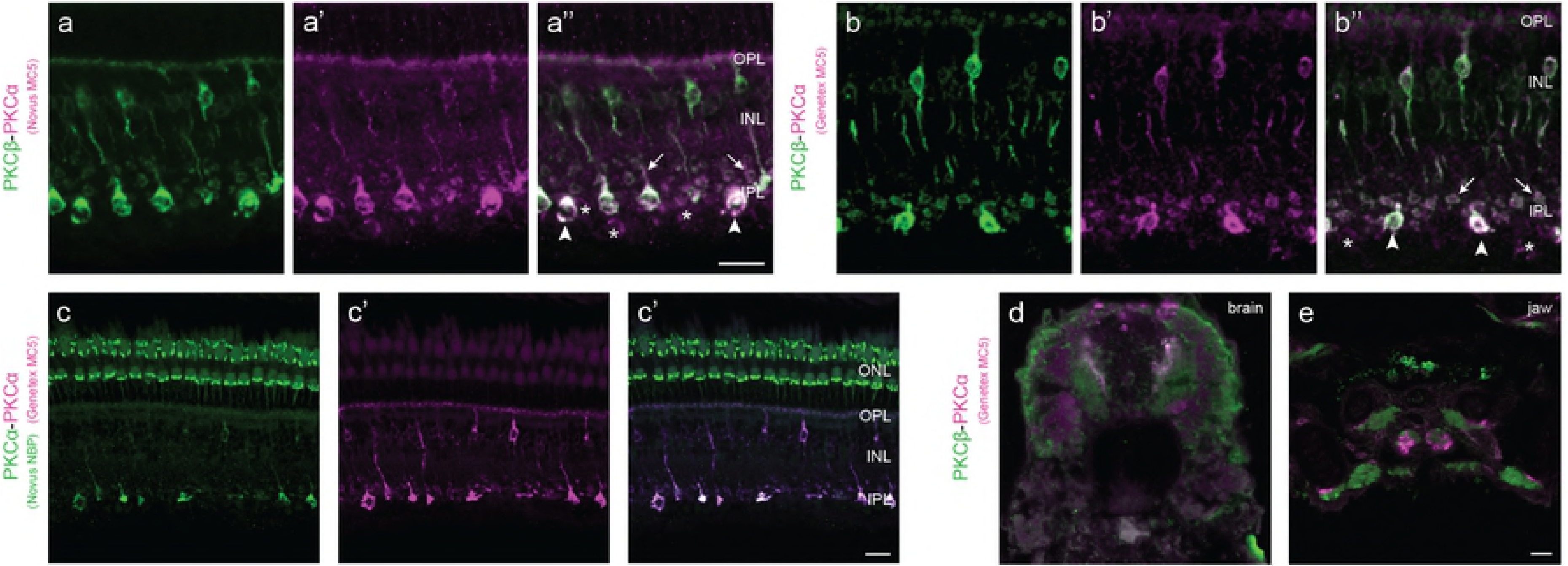
Commercially available PKC antibodies label overlapping but also distinct central nervous system structures. Double labeling of PKCα (MC5 Novus, MC5 Genetex, and NBP Novus) and PKCβ (Santa Cruz Biotechnology) antibodies on retinal sections of adult (a-c) and 5 dpf larval (d,e) zebrafish. Both PKCα MC5 antibodies show an overlapping labeling in retinal bipolar cells with the PKCβ antibody (a” and b”). Different shapes of axon terminals in the IPL (arrows and arrowheads in a” and b”) suggest labeling of at least two different bipolar cells types. In addition, both PKCα MC5 antibodies weakly label some cells in the GCL (asterisk in a” and b”). The epitope of a third PKCα antibody (NBP, Novus) is located in an overlapping manner in the INL and IPL but is also found accessory outer segments of photoreceptors in the ONL (c-c”). Double labeling on transverse sections of 5 dpf larvae shows that the PKCα (MC5, Genetex) and –β antibodies additionally label different areas of the brain and the jaw (d,e). For abbreviations, see list. Scale bar in a” (applies to a’-” and b’-”), in c” (applies to c’-”) and in e (applies to d and e) = 10 μm.

### Expression of recombinant PKCs and Western blot

The coding region of each zebrafish *prkc* was PCR amplified with the primers that are listed in Table 1, and subcloned into pIRES2:EGFP vector (kindly provided by M. Kamermans). Expression was done in HEK293T cells. Transient transfection of cells using the Ca_3_(PO_4_)_2_ technique was performed as previously described (Kimmel, Ballard, Kimmel, Ullmann, & Schilling, 1995). 5 μg linearized DNA or only buffer for mock transfection was added to the cells. 30 to 40 hours post-transfection when the cell layer had reached a confluence of up to 100%, cells were checked for GFP signals. Subsequently, the medium was aspirated and the plates were placed on ice and washed with three times with 3 ml cold PBS containing 0.9 mM CaCl_2_. Next, cells were lysed by adding 500 μl Laemmli buffer (4.4 ml 0.5M Tris-HCl pH 6.8, 4.4 ml Glycerol, 2.2 ml 20% SDS, 0.65 ml 1% Bromophenol blue) supplemented with Protease inhibitor (Complete Mini, Roche), and collected in a 2 ml Eppendorf tube. Cell lysates were supplemented with 1:40 β-Mercaptoethanol and homogenized with a pistil. After heating them to 90°C for 5 min, the lysates were cleared using centrifugation, the supernatant sonicated and subjected to Western blot analysis. Lysates were loaded on a 10% precast gel (Mini Protean TGX, Biorad, Cressier, Switzerland), blotted to PVDF membranes (0.2 μm, Novex, Thermo Fisher Scientific) which were blocked for 2 hours in PBS containing 0.05% Tween 20 and 3% dry milk powder (PBS-TM) at RT. Primary antibodies were used in the same concentrations as described for immunohistochemistry and applied ON at 4°C in PBS-TM. As a loading control anti-Vinculin (124 kDa; 1:5000, Genetex) was used. After a 5 min washing step in PBS-TM followed by two 10 min washing steps in PBS containing 0.05% Tween 20 (PBT), the membranes were incubated for 45 min at RT with secondary horseradish peroxidase (HRP-) linked antibodies (Invitrogen, Thermo Fisher Scientific) diluted in PBS-TM (goat anti-rabbit, 1:5000; goat anti-mouse, 1:7500). Following a 20 min washing with PBS-TM and four 5 min washes with PBT, membranes were subjected to development solution (Super Signal West Dura Extended Duration Substrate, Thermo Fisher Scientific) for 5 min at RT. Finally, the signals were detected by the LAS 4000 Chemiluminescence Imager (software: Image Quant LAS 4000, automatic exposure) and processed using Adobe Illustrator C5.

**Table 1:**
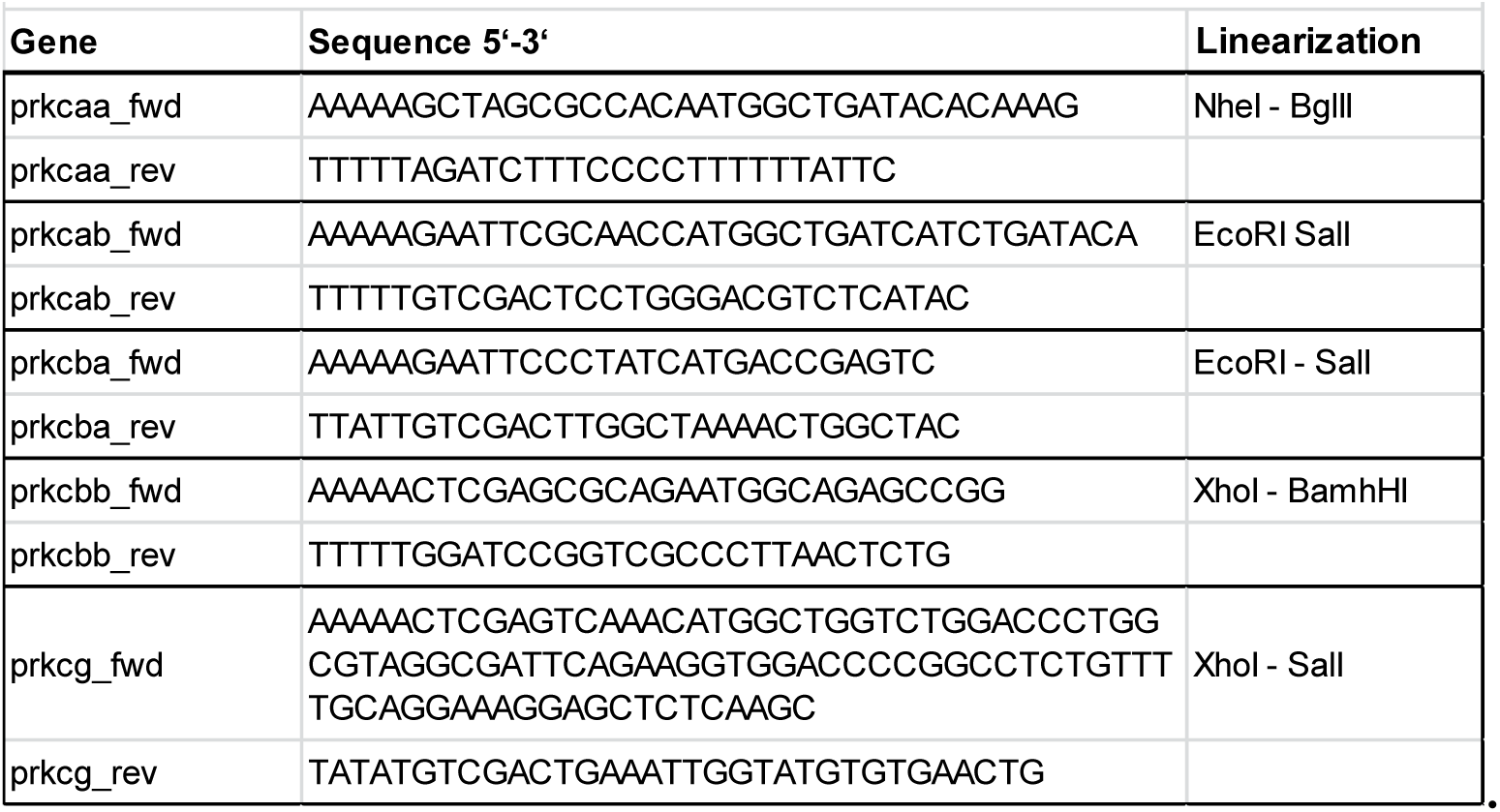
Primerss used for the cloning of expression constructs.

**Table 2:**
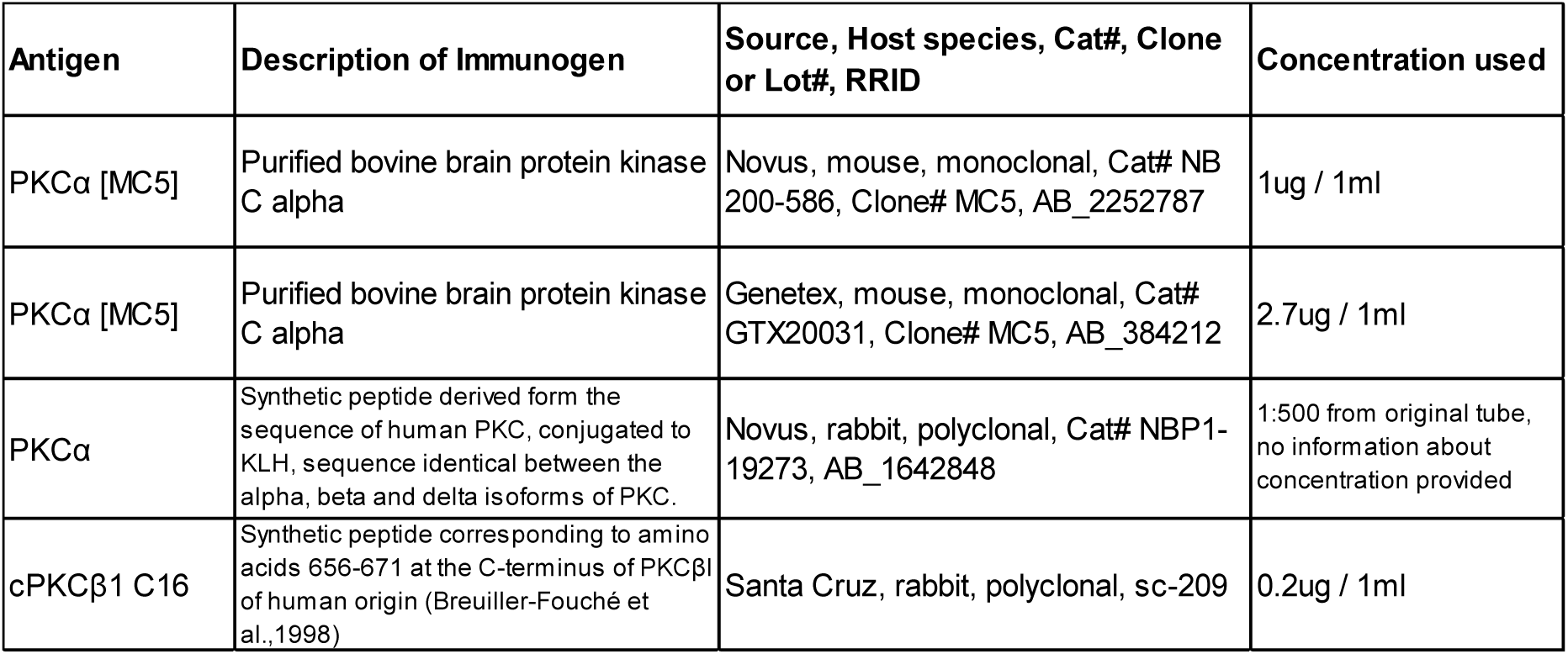
Overview of the antibodies used in this study.

## Results

In mammals, the family of conventional *prkcs* consists of the three members -*a*, -*b* and -*g* (Newton, 2010). Based on sequence similarity, we annotated and cloned five different zebrafish *prkc* cDNAs, two paralogs of *prkca* and *prkcb* and one single *prkcg* paralog. The phylogeny and the detailed description of the larval expression pattern of these genes is reported in [35]. In this study we describe *prkc* transcript expression in the retina by *in situ* hybridization in combination with PKC antibody labeling using commercially available antibodies.

### Commercially available PKCα and -β antibodies label overlapping subsets of ON-bipolar cells

PKCα and -β antibodies are used as a marker for retinal ON-bipolar cells in different species [2,36,12]. As there are no zebrafish specific PKC antibodies available commercial PKC antibodies are commonly used as bipolar markers. We tested different frequently used PKC antibodies raised against bovine and human epitopes and found marked differences in their staining profile of bipolar cells (Fig 1).

All used PKC antibodies showed specific labeling in the retinal INL and IPL, presumably in ON-bipolar cells and their processes (Fig 1A-C). A separate double labeling of each PKCα MC5 with PKCβ showed that all antibodies label identical cells in the middle part of the INL (Fig 1A’-” and B’-”), some with smaller axon terminals (arrows) and some with a larger axon terminal (arrowheads). In addition, both PKCα MC5 antibodies detect cells in the GCL (asterisk in Fig 1A”,B”,3J). Another PKCα antibody (Novus (NBP)) also weakly labels the same cells in the INL but showed a very strong labeling in the retinal ONL, presumably in accessory outer segments (Fig 1C’-”) (Hodel et al., 2014). Aside from the retina, PKC antibodies additionally label different cells in other tissues. Applying PKCα MC5 (Genetex) and PKCβ on transverse sections of the brain and the jaw of 5 days old zebrafish larvae shows antibody-specific labeling in distinct areas of both examined tissue samples (Fig 1D,E), demonstrating that these antibodies recognize different zebrafish PKCs with different affinities. Hence, the labeled ON-bipolar cells might express a mix of different PKCs.

### *prkc* transcripts in the zebrafish retina are expressed in overlapping but distinct patterns

To gain a detailed view of *prkc* expression in the zebrafish retina, we analyzed adult retinal tissue by *in situ* hybridization (ISH). While both paralogs of the zebrafish *prkca* and –*b* genes are expressed in the adult zebrafish retina, we never observed expression of *prkcg* (Fig 2a,c,e,g,). Therefore, we excluded *prkcg* from further analysis. In contrast to *prkcaa* mRNA that can be detected in the middle INL (Fig 2a), *prkcab* and the two *prkcb* transcripts are more widely expressed. *prkcab* is expressed in the middle and the distal INL, as well as in the GCL (Fig 2c). A strong labeling in the proximal INL and the GCL is seen for *prkcba* (Fig 2e) whereas *prkcbb* is only weakly expressed throughout the INL and in the GCL (Fig 2g). The expression pattern we found using the same probe but with a fluorescent tag (FISH) were generally overlapping (Fig 2 b1,d1,f1,h1). However, for *prkcbb* we found an additional weak expression in the ONL (arrowhead in Fig 2h1), suggesting a difference in the sensitivity of the two detection methods.

**Figure 2:**
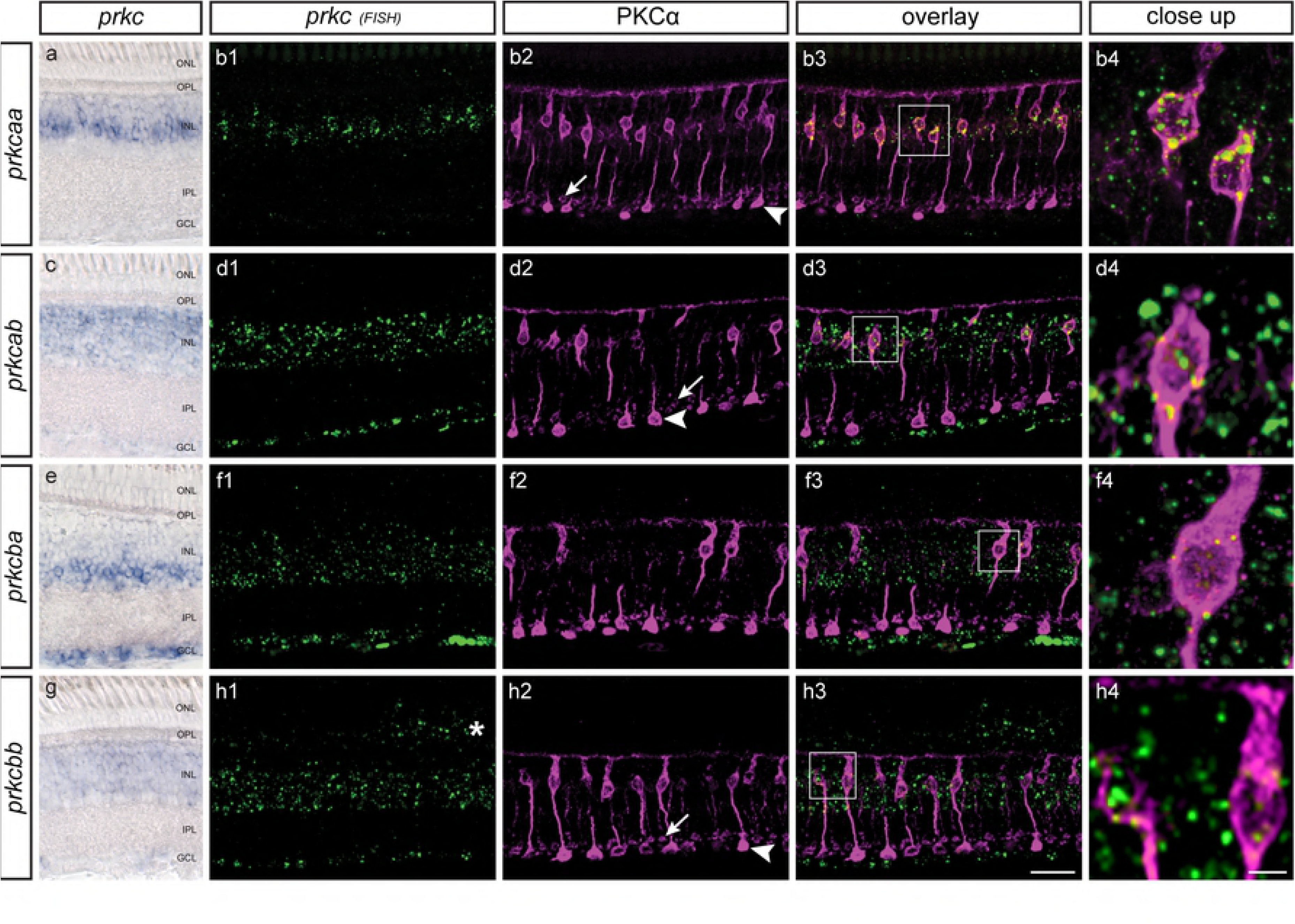
Co-expression analysis of *prkca* and –*b* transcripts with PKCα antibody labeling in adult retinal sections. Conventional (blue) and fluorescent (green) mRNA labeling of *prkca* and –*b* paralogs (a,c,e,g and b1,d1,f1,h1). Both *prkca* paralogs are expressed in the inner nuclear layer (INL) of the retina. While the expression of*prkcaa* is restricted to the middle INL (a, b1),*prkcab* transcripts show a broader expression in the middle and the distal INL, and in the ganglion cell layer (GCL; c, d1). *prkcba* is expressed in the proximal INL and strongly in the GCL (e, f1), and its paralog, *prkcbb*, weakly throughout the INL and GCL (g, h1). Additionally photoreceptors are weakly labeled by the fluorescent method (asterisk in h1). Confocal images of adult retinal sections showing fluorescent *in situ* hybridization (green) in combination with PKCα MC5 (Novus) antibody labeling (magenta) and the corresponding overlay (b3,d3,f3,h3). PKCα labeling is found in two different types of bipolar cells, as illustrated with arrows vs. arrowheads. We find expression of *prkcaa* in all PKCα-positive bipolar cell bodies in the middle INL (b3-4), while *prkcab* and –*bb* show only partial overlap (d3-4, h3-4). *prkcba* transcripts seem not to overlay with PKCα-positive cells (f3-4). For abbreviations, see list. Scale bar in h3 (applies to all images except the close ups) = 25 μm. Scale bar in h4 (applies to b4,d4,f4,h4) = 5 μm.

### PKCα MC5 antibody labeling highlights prkcαa, –αb, and –βb expressing cells and the corresponding proteins

As the different zebrafish *prkcs* were not expressed in the same retina layers, we combined *prkc* RNA labeling with antibody staining in adult retinal sections to demonstrate which *prkc* expression overlaps with the PKC antibody labeling. We chose to use PKCα MC5 of Novus as a marker for this experiment, as it shows an overlapping labeling in bipolar cells with all other antibodies tested but also labeling in some additional cells in the GCL compared to PKCβ (Fig 1a”).

Interestingly, all PKCα-positive bipolar cells clearly express *prkcaa* within their cell bodies and vice versa (Fig 1b3-4), whereas the *prkcab* transcript seems to be located in some but not all bipolar cells labeled by the PKCα antibody (Fig 1d3-4). For *prkcb* genes we found no overlap of the antibody labeling with *prkcba* (Fig 1f3-4) and only a partial overlap with *prkcbb* (Fig 1h3-4).

In order to gain insight into PKC antibody specificity in zebrafish we generated expression constructs of full length *prkc* transcripts, and tested antibody recognition of recombinant proteins by Western blot (Fig 3). Since both PKCα MC5 antibodies showed comparable results, only the result with the PKCα MC5 from Genetex is shown. For each antibody a different pattern can be observed (see overview in Fig 3d). When applying the PKCα NBP antibody on recombinant zebrafish PKCs, bands in different intensities around the expected size of 75 kDa can be detected (Fig 3a). The PKCα NBP antibody recognizes a faint band of a lower size for PKCβa (black arrowhead in Fig 3a, 3. lane) and a strong band at a higher position for PKCβb (white arrowhead in Fig 3a, 4. lane). The PKCα MC5 antibody recognizes the zebrafish PKCαb at exactly 75 kDa (Fig 3b, 2. lane), and in addition PKCαa and βKCβb at a slightly higher position (Fig 4b, 1. and 4. lane. The PKCβ1 antibody strongly recognizes recombinant zebrafish PKCαa (Fig 3c, 1. lane) and weakly PKCβb (Fig 3c, 4. Lane), but none of the other recombinant PKCs (Fig 3c, 2., 3., 5. and 6. lane). These western blot results confirm that these antibodies recognize various antigens, explaining their differential immunohistological labeling of differing bipolar cell populations.

**Figure 3:**
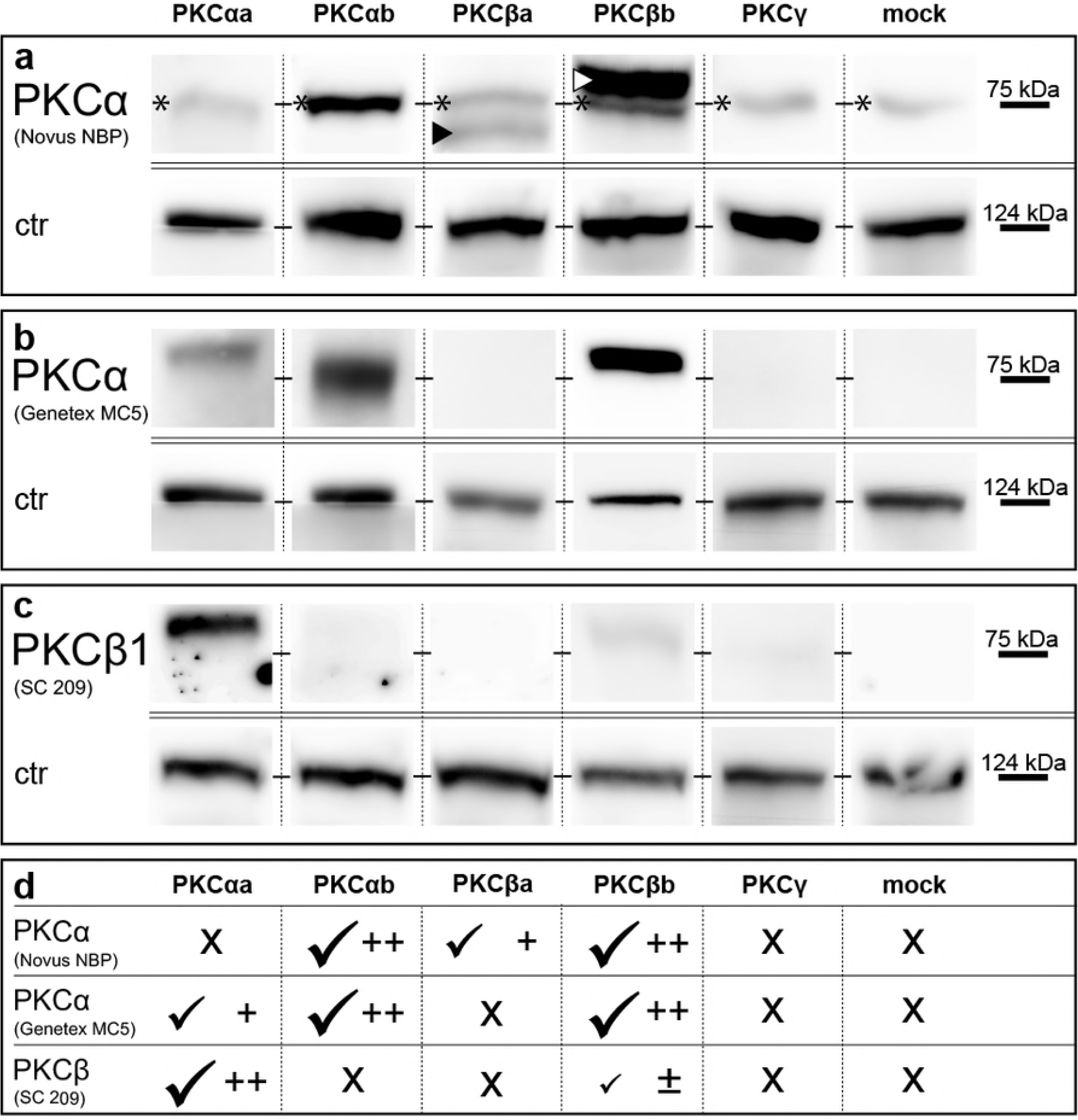
Western blot analysis of recombinant zebrafish cPKC proteins using PKCα and –β antibodies. Three different antibodies were used to compare the specificity of each antibody with recombinant zebrafish PKCs. The PKCα NBP antibody is specific for PKCαb and both PKCβ paralogs. Besides, PKCα NBP detects a faint band in all samples, including mock controls, likely recognizing an endogenous distantly related PKC orthologue present in HEK293T cells (asterisk at 75 kDa), PKCα MC5 recognizes both zebrafish PKCα paralogs (b, 1. and 2. lane) as well as PKCβb (b, 4. lane). The PKCβ1 antibody also recognizes PKCαa (c, 1. lane) but PKCβb only faintly (c, 4. lane), and PKCαb not at all (c, 2. lane). PKC*γ* protein was not specifically detected with any antibody used (a-c, 5. lane). Anti-Vinculin (124 kDa) antibodies were used as loading controls. The different antibody recognition patterns are summarized in d.

## Discussion

After analyzing the expression and phylogenetic relations of zebrafish *prkc* genes (Haug, Gesemann, Berger, & Neuhauss, 2018) we focused on the retina, as PKCα and -β antibodies are widely used in the community as markers for ON-bipolar cells. To identify which zebrafish PKC(s) are labeled by the commercially available antibodies we performed fluorescent ISH in combination with PKCα antibody labeling and Western blots using recombinant PKCs.

### Commercially available PKC antibodies recognize different zebrafish PKC variants

Although non-mammalian vertebrates possess a significantly higher diversity of bipolar cells than mammals, their main functions are conserved [37]. PKCα and sometimes PKCβ antibodies are commonly used as ON-bipolar cell markers. However, the frequently used antibodies against PKCα and/or -β have been designed to recognize human or bovine epitopes. Hence, it is not known which PKC proteins are recognized in other species. Moreover, the whole genome duplication event in the lineage of teleost fish has added to the complexity [34], increasing the number of *prkcs* to five. This increase in retinal genes may be the basis for the 33 different types of bipolar cells that were recently identified in zebrafish [20]. Another explanation could be that some aspects of visual processing that takes place in higher brain areas of mammals is achieved in the retina of lower vertebrates.

Our initial ISH showed labeling of both *prkca* and *prkcb* paralogs in the INL of the retina where bipolar cells are located. We found a complete overlap in cells of the middle INL when applying the riboprobe of *prkcaa* and the antibody against PKCα MC5 (Fig 2b1-4). Some but not all PKCα-positive cell bodies did also express *prkcab* and *prkcbb* (Fig 2d1-4, h1-4), indicating that at least some PKCα-positive cells express different *prkc* paralogs, while others only seem to exclusively express *prkcaa.* Interestingly, while the PKCα MC5 antibody indeed recognizes the zebrafish PKCαa, it also recognizes other zebrafish PKCs (Fig 3). Moreover, western blot results indicated that all tested PKC antibodies recognize a combination of PKCs, demonstrating that the tested antibodies are not specific for one single zebrafish PKC but rather label a mixture of different PKC subtypes. When tested on other tissues such as the brain and the jaw of 5 day old larvae, the PKCα MC5 and the PKCβ antibodies were expressed in clearly different areas, indicating that these two antibodies indeed do not recognize the same combination of zebrafish PKCs as also shown by Western blots (Fig 3).

### PKCα and PKCβ antibodies as marker for ON-bipolar cells

It is generally assumed that PKCα [10] or PKCα/β antibodies [12] label B_ON_ s6L cells, a bipolar cell type morphologically resembling the mixed-input (b1 or Mb) ON-bipolar cells of other teleosts [19]. Recent data suggest that the B_ON_ s6L bipolar cell is identical to the RRod cell, an ON-bipolar cell that only contacts rods [20]. However, earlier studies describe labelling of an additional bipolar cell type by PKCα or PKCα/β antibodies [38,12], which possibly corresponds to the slightly smaller B_ON_ s6 type that contacts cones [19].

All tested antibodies labeled the same cells of the INL in adult zebrafish retina and they seem to be of at least two kinds: we find labeled cells that contain large axon terminals in a more proximal part of the IPL as well as other cells with smaller axon terminals that ramify in a more distal part of the IPL. Based on morphology, these two subtypes likely comprise the above mentioned B_ON_ s6L or RRod type and the B_ON_ s6 type [19,20]. As *prkcaa* expression and the antibody labeling are identical, this indicates that *prkcaa* is expressed in these two different bipolar cell subtypes as well.

The second *prkca* paralog, *prkcab*, is also expressed in the middle INL, however its expression only partially overlaps with PKCα-positive cells. This different expression suggests a gain of function after the teleost-specific whole genome duplication [39,34]. Interestingly, *prkcbb* is also located in the middle INL and overlaps with some PKCα-positive cells, however, it might also label additional subtypes.

Intriguingly, in larval tissue both *prkca* paralogs label cells in the middle INL [35] but expression in other retinal cells is only visible in adult sections (Fig 1). In addition, both *prkcb* paralogs are only expressed in the adult retina (Fig 1) [35]. For most species it is hypothesized that PKCα and/or -β labels additionally rod ON-bipolar cells [40,4,1,7] and our study also indicates the labeling of RRod cells. As the rod circuitry is established only at later stages and has been shown to be functional earliest in 15 day old larvae [41] expression of molecules involved in rod ON-bipolar cell signaling are expected to appear only in older larvae. Taken together our results suggest that PKC variants are also expressed in RRod cells and that the commercial antibodies recognize at least a subset of these cells.

### *prkc* expression in other retina cells

Besides the expected expression in bipolar cells, both *prkca* and -*b* paralogs were also found in additional retinal cell types. So far, studies describe the expression of conventional PKCs in photoreceptors [42–47], and in rod outer segments [48,49]. However, other studies were unable to detect PKC-positive photoreceptors (e.g. [4,50,1] or show ambiguous results depending on the applied technique [51,52] or the species examined [10,7]. While we did not detect any *prkc* transcripts in photoreceptors by conventional ISH, fluorescent ISH of *prkcbb* showed a weak expression in the ONL suggesting that low concentration of transcripts are indeed present. This is in line with reports about the crucial role of PKCs in photoreceptor development [53] as well as phosphorylation of a number of molecules important for phototransduction (e.g. rhodopsin) [54,49,55]. Interestingly, the PKCα NBP antibody (Fig 1C’,C”) shows a similar expression in the photoreceptor layer as was previously published by Osborne and colleagues for a PKCα labeling in the rabbit retina (Osborne, Barnett, Morris, & Huang, 1992). As the company states on the product datasheet that this antibody recognizes the PKCα, –β, and –δ isoform, the labeling in photoreceptors might be due to any of these PKCs.

*Prkcab*, -*ba* and -*bb* all show an additional expression in the INL besides the expected expression in bipolar cells. Due to the location in the distal or proximal INL, respectively, we expect *prkcab* to be expressed in horizontal cells and *prkcba* in amacrine cells while *prkcbb* is distributed throughout the INL and might be expressed in both retinal cell types. Earlier investigations have shown an involvement of PKCs in activity dependent morphological changes of horizontal cell synapses [56,57], however, this was disputed later [58]. Vertebrate amacrine cells, however, do definitely also express conventional PKCs. This was shown by subtype-specific labeling of PKCα in rat and rabbit [51,47], and PKCβ in rat and human [50,51]. As we only found evidence for *prkcba* and maybe *prkcbb* expression in the proximal INL where amacrine cells are located, one of the PKCβ-paralogs might has cover the function of the PKCα in these cells in zebrafish.

Interestingly, at least in the rodent retina PKC *γ* has also a function, as the lack of PKC *γ* (but also of PKCβI) totally inhibits rod development in mice [53]. Previous studies are conflicting with some reporting expression of PKCγ in the retina [50,45], while others failed to detect any labeling [43,51,47,59]. In our study the absence of retinal expression of *prkcg* cannot be attributed to technical issues, since the *prkcg* probe shows distinct cerebellar expression [35].

We find a weak *prkcab* and –*bb* expression as well as a very pronounced expression of *prkcba* in the retinal ganglion cell layer of adult zebrafish, however, in an earlier study we did not detect any *prkc* expression in the GCL in larval tissue [35]. This is in accordance with previous studies, where PKCβ has been located in the GCL [43,50,51,45,47] but only few studies describe the expression of PKCα in this retinal layer [60,59]. In line with the transcript expression, we find the PKCα MC5 antibody of both companies to label cells in the GCL (asterisk in Fig 1A”,B”, 3J), indicating that these antibodies recognize at least one of the β-paralogs as well.

## Conclusion

Commercial PKCα and –β antibodies are commonly used to label ON-bipolar cells in the vertebrate retina. We found that these antibodies indeed consistently label a subset of ON-bipolar cells of both the scotopic and photopic pathway (B_ON_ s6L or RRod type and B_ON_ s6) in the zebrafish retina. However, these antibodies are not as often considered pan-ON-bipolar cell markers but rather mark different bipolar cell subtype populations.

## Acknowledgements

We would like to acknowledge Kara Kristiansen and Martin Walther for technical support and excellent fish care.

